# Climate and seed weight drive intraspecific variation in seed longevity in storage

**DOI:** 10.1101/2025.01.27.635048

**Authors:** Lea Klepka, Sascha Liepelt, Anna Bucharova

## Abstract

**Premise:** Conservation seed banks are essential for ex-situ plant conservation, but stored seeds slowly deteriorate and lose viability. Seed longevity in storage is determined by the initial seed viability and the rate of seed viability loss. The rate of seed viability loss in storage varies between species, and there is possibly also some variation between populations or even genotypes within species. However, the extent of this intraspecific variability and its drivers remain unclear.

**Methods:** We investigated both inter- and intraspecific variability in seed longevity and its predictors in 41 common grassland species and 188 seed accessions from across Europe. We exposed the seeds to artificial ageing conditions (60% RH, 45°C) and used probit analysis to obtain the rate of seed viability loss (σ) as a measure of seed longevity. We then related σ to both accession- and species-specific factors.

**Key results:** Seed longevity (σ) varied significantly among accessions within 58% of the species, and the probability of detecting such intraspecific differences increased with the number of accessions available for a given species. This suggests that within-species variation in seed longevity is widespread. Accession-specific predictors explained only 14.4% of the within-species variability. Specifically, seed longevity increased with the mean annual temperature at the accession origin and decreased with the accession-specific seed weight. Across species, seed longevity differed among plant families but was unrelated to seed weight or seed chemical composition.

**Conclusions:** Our findings highlight substantial within-species variation in seed longevity in storage, which, however, is difficult to predict.

## INTRODUCTION

Biodiversity conservation is one of the critical global challenges, and *ex situ* methods play an increasingly important role in this task (Cohen et al., 1991). One of the most common ex-situ methods for plant conservation is seed banks, dedicated facilities that store seeds of wild plant species, particularly the vulnerable ones (Peres, 2016). The stored seeds represent a snapshot of past populations and are valuable resources for species reintroductions or ecosystem restoration, and also for scientific research (Cochrane et al., 2007; Etterson et al., 2016; Wambugu et al., 2023). Thus, the ultimate task of conservation seed banks is to keep the seeds viable for decades or even centuries.

Seed survival in storage depends on the type of seeds and storage conditions. Some types of seeds, so-called recalcitrant seeds, do not survive desiccation, making long-term storage impossible. Other types of seeds, called orthodox, can survive desiccation, and they can be stored for a couple of months up to centuries, depending on the species (Hay & Probert, 2013). Conservation seed banks aim to maintain seed viability as long as possible by storing the seeds under low temperatures and humidity to slow down internal chemical processes such as lipid peroxidation and DNA damage (Nagel et al., 2015; Walters et al., 2005a). The seed longevity also depends on the quality and viability of the seeds (Hay & Probert, 2013; Niedzielski et al., 2009), and thus, guidelines for seed banking recommend storing accessions with high initial viability (ENSCONET, 2009; Millennium Seed Bank, 2022). Yet, even highly viable seeds stored under optimal conditions slowly deteriorate and eventually die. This mortality is faster when the seeds are stored in suboptimal conditions, for example when storing seeds for large-scale ecosystem restoration in ambient conditions at a farm (Merritt & Dixon, 2011). To optimise the long-term effectiveness of seed storage – be it for conservation, research or ecosystem restoration purposes – we need to understand which factors predict seed longevity in storage.

Studying seed longevity presents unique challenges, as it requires comparing old and fresh seeds of the same seed lot, which would need time travel. To circumvent this obstacle, researchers have developed artificial ageing protocols which simulate natural ageing by exposing the seeds to high temperatures and high relative humidity (Delouche & Baskin, 2021). These conditions accelerate the same or at least similar chemical processes typical of natural ageing during long-term storage (Delouche & Baskin, 2021). Although this method may not perfectly replicate natural ageing, the longevity of seeds in artificial ageing correlates with the longevity of seeds stored in seed banks both across and within species (Ninoles et al., 2022; Probert et al., 2009). Nevertheless, artificial ageing is the best method we currently have to compare seed longevity among accessions.

Seed survival in artificial ageing differs between species, and to some degree also between higher taxa like plant families, although large plant families harbour substantial variation in seed longevity (Probert et al., 2009; Walters et al., 2005a). Studies aiming to detect general patterns in seed longevity beyond evolutionary history yielded mixed results. Endospermic seeds survive shorter in storage than non-endospermic ones (Probert et al., 2009). The influence of seed weight is less clear; some studies suggest that seeds of small-seeded species survive longer (Satyanti et al., 2018), while others report no effect of seed weight (Davies et al., 2020; Merritt et al., 2014; Probert et al., 2009). Seed longevity is also affected by climatic conditions at seed collection sites: Seeds from hot or dry environments generally survive longer than those from cooler or wetter conditions (Merritt et al., 2014; Probert et al., 2009; Zani & Müller, 2017). Furthermore, seed longevity should hypothetically depend on the seed’s chemical composition because high oil content should reduce seed longevity due to ongoing lipid peroxidation in storage (Narayana Murthy & Sun, 2000; Pritchard & Dickie, 2003), yet the results of empirical studies are inconsistent (Nagel et al., 2015; Probert et al., 2009; Walters et al., 2005a).

In contrast to between-species comparison, there is less comprehensive information on within-species variability on seed longevity in storage. Most data come from cultivated crops or model species (e.g. Bizouerne et al., 2023; Franks et al., 2018; Guzzon et al., 2021; Lee et al., 2019), with only a few studies addressing natural populations of wild species (Genna et al., 2020; Kochanek et al., 2009; Mondoni et al., 2011; White et al., 2023). These indicate that the variation in seed longevity within a species can be as substantial as the variation between species from the same ecosystem (Kochanek et al., 2009; Mondoni et al., 2011). The predictors of seed longevity on the level of populations, genotypes, or varieties appear to be similar to those driving between-species differences: In general, lighter seeds of the same species typically survive longer than heavier seeds (Franks et al., 2019; Guzzon et al., 2018; Lee et al., 2019, but see Mira et al., (2019)). Consequently, seeds of genotypes or varieties with lighter seeds tend to survive longer in storage than accessions with heavier seeds, although this effect is inconsistent across studies (Guzzon et al., 2021; Schutte et al., 2008). The climate at the seeds’ origin seems to be also a critical factor, with populations from warmer and dryer environments producing longer-lived seeds than populations from colder and more humid environments (Mondoni et al., 2011, 2014, but see Kochanek et al. (2009)). The differentiation in seed longevity among populations of the same species can be driven by two, mutually not exclusive mechanisms: genetic differentiation (Mondoni et al., 2014) or phenotypic plasticity, where parental growing conditions influence seed longevity (Kochanek et al., 2010; Mondoni et al., 2014; Sinniah et al., 1998). While the studies above provide valuable insights, we are still missing a comprehensive assessment of intraspecific variability in seed longevity across wild species, its magnitudes and predictors.

To fill this gap, we focused on more than 40 common grassland species represented by up to 11 accessions from across Europe. We exposed the seeds to artificial ageing, estimated their longevity, and analysed the data on two levels: Comparison across accessions within the species and comparison across species.

We expect that (1) seed longevity under artificial ageing conditions differs between species and accessions within species. The variation among species will be larger than the variation within species. (2) Within species, longevity will increase with mean annual temperature and annual precipitation at the seed origin and decrease with seed mass. (3) Across species, longevity will decrease with seed weight and be influenced by seed chemical composition, particularly oil content, and the plant family.

## MATERIALS AND METHODS

### Study species and seed material

We focused on 41 species common in European grasslands, belonging to 11 families distributed across the phylogenetic tree of angiosperms (Table 1). They represent a wide variety of traits, some potentially associated with seed longevity, specifically seed weight, seed oil and protein content (Probert et al., 2009). All species have orthodox seeds, i.e. they survive desiccation and freezing during storage.

**Table 1:**
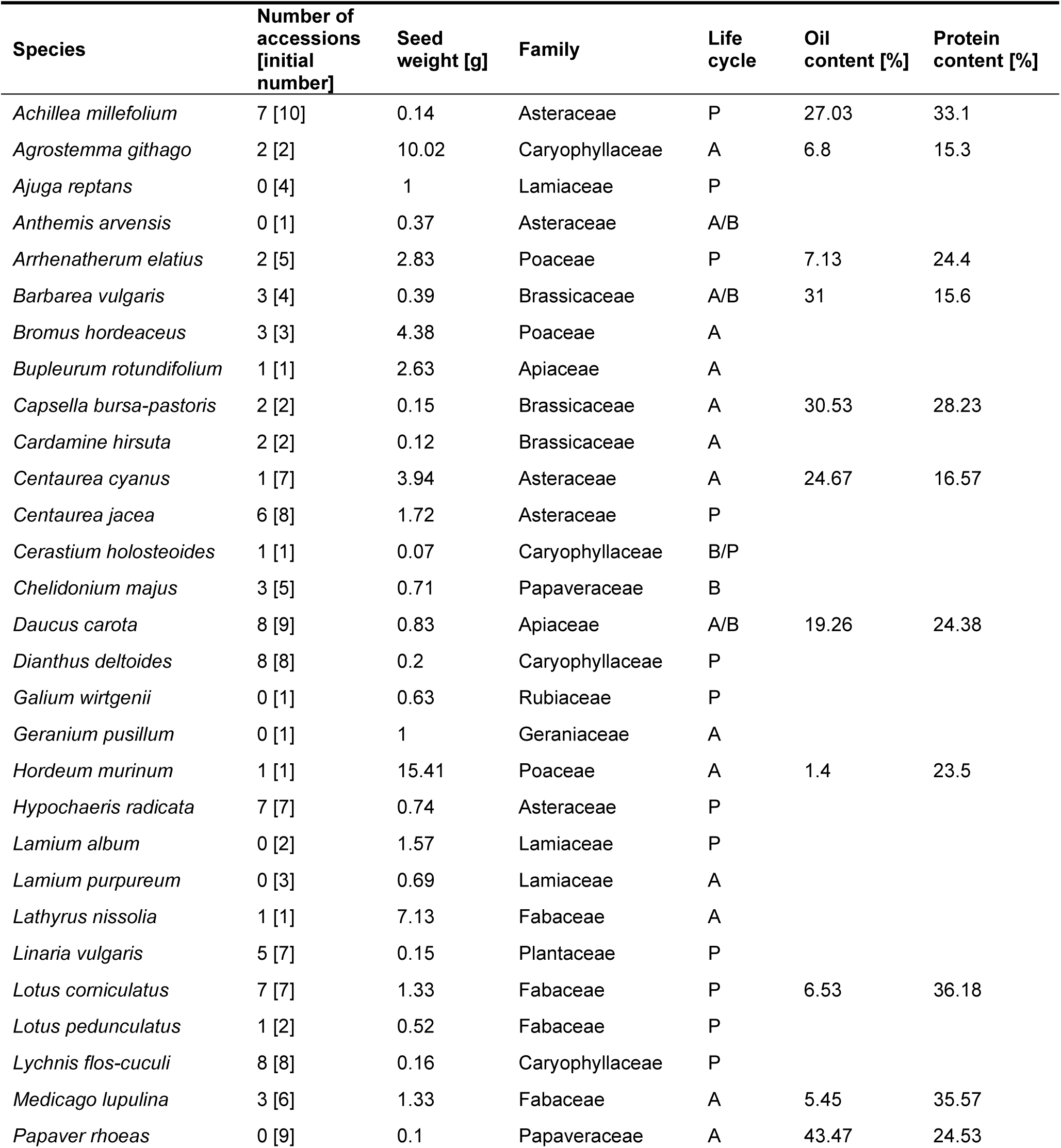
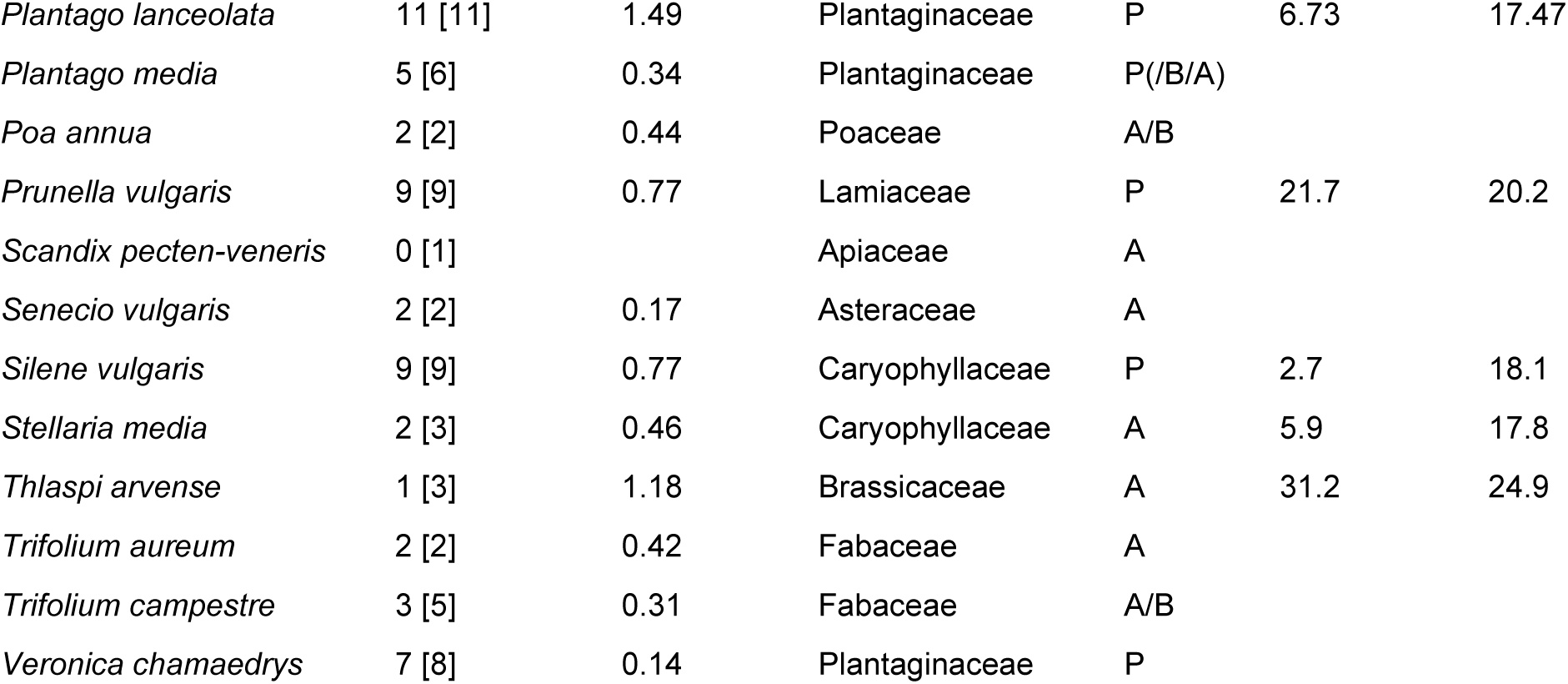
Species included in the study. The number of accessions is the number of accessions used for the analyses, and in brackets is the initial number of accessions. Seed weight is the weight of 1000 seeds, averaged across all accessions of the same species. Oil and protein content are obtained from the Seed Information Database (Society for Ecological Restoration et al., 2023). Life cycle: A = annual, P = perennial, B = biennial. *: Information of conspecific

| Species | Number of accessions [initial number] | Seed weight [g] | Family | Life cycle | Oil content [%] | Protein content [%] |
| --- | --- | --- | --- | --- | --- | --- |
| *Achillea millefolium* | 7 [10] | 0.14 | Asteraceae | P | 27.03 | 33.1 |
| *Agrostemma githago* | 2 [2] | 10.02 | Caryophyllaceae | A | 6.8 | 15.3 |
| *Ajuga reptans* | 0 [4] | 1 | Lamiaceae | P |  |  |
| *Anthemis arvensis* | 0 [1] | 0.37 | Asteraceae | A/B |  |  |
| *Arrhenatherum elatius* | 2 [5] | 2.83 | Poaceae | P | 7.13 | 24.4 |
| *Barbarea vulgaris* | 3 [4] | 0.39 | Brassicaceae | A/B | 31 | 15.6 |
| *Bromus hordeaceus* | 3 [3] | 4.38 | Poaceae | A |  |  |
| *Bupleurum rotundifolium* | 1 [1] | 2.63 | Apiaceae | A |  |  |
| *Capsella bursa-pastoris* | 2 [2] | 0.15 | Brassicaceae | A | 30.53 | 28.23 |
| *Cardamine hirsuta* | 2 [2] | 0.12 | Brassicaceae | A |  |  |
| *Centaurea cyanus* | 1 [7] | 3.94 | Asteraceae | A | 24.67 | 16.57 |
| *Centaurea jacea* | 6 [8] | 1.72 | Asteraceae | P |  |  |
| *Cerastium holosteoides* | 1 [1] | 0.07 | Caryophyllaceae | B/P |  |  |
| *Chelidonium majus* | 3 [5] | 0.71 | Papaveraceae | B |  |  |
| *Daucus carota* | 8 [9] | 0.83 | Apiaceae | A/B | 19.26 | 24.38 |
| *Dianthus deltoides* | 8 [8] | 0.2 | Caryophyllaceae | P |  |  |
| *Galium wirtgenii* | 0 [1] | 0.63 | Rubiaceae | P |  |  |
| *Geranium pusillum* | 0 [1] | 1 | Geraniaceae | A |  |  |
| *Hordeum murinum* | 1 [1] | 15.41 | Poaceae | A | 1.4 | 23.5 |
| *Hypochaeris radicata* | 7 [7] | 0.74 | Asteraceae | P |  |  |
| *Lamium album* | 0 [2] | 1.57 | Lamiaceae | P |  |  |
| *Lamium purpureum* | 0 [3] | 0.69 | Lamiaceae | A |  |  |
| *Lathyrus nissolia* | 1 [1] | 7.13 | Fabaceae | A |  |  |
| *Linaria vulgaris* | 5 [7] | 0.15 | Plantaceae | P |  |  |
| *Lotus corniculatus* | 7 [7] | 1.33 | Fabaceae | P | 6.53 | 36.18 |
| *Lotus pedunculatus* | 1 [2] | 0.52 | Fabaceae | P |  |  |
| *Lychnis flos-cuculi* | 8 [8] | 0.16 | Caryophyllaceae | P |  |  |
| *Medicago lupulina* | 3 [6] | 1.33 | Fabaceae | A | 5.45 | 35.57 |
| *Papaver rhoeas* | 0 [9] | 0.1 | Papaveraceae | A | 43.47 | 24.53 |
| *Plantago lanceolata* | 11 [11] | 1.49 | Plantaginaceae | P | 6.73 | 17.47 |
| *Plantago media* | 5 [6] | 0.34 | Plantaginaceae | P(/B/A) |  |  |
| *Poa annua* | 2 [2] | 0.44 | Poaceae | A/B |  |  |
| *Prunella vulgaris* | 9 [9] | 0.77 | Lamiaceae | P | 21.7 | 20.2 |
| *Scandix pecten-veneris* | 0 [1] |  | Apiaceae | A |  |  |
| *Senecio vulgaris* | 2 [2] | 0.17 | Asteraceae | A |  |  |
| *Silene vulgaris* | 9 [9] | 0.77 | Caryophyllaceae | P | 2.7 | 18.1 |
| *Stellaria media* | 2 [3] | 0.46 | Caryophyllaceae | A | 5.9 | 17.8 |
| *Thlaspi arvense* | 1 [3] | 1.18 | Brassicaceae | A | 31.2 | 24.9 |
| *Trifolium aureum* | 2 [2] | 0.42 | Fabaceae | A |  |  |
| *Trifolium campestre* | 3 [5] | 0.31 | Fabaceae | A/B |  |  |
| *Veronica chamaedrys* | 7 [8] | 0.14 | Plantaginaceae | P |  |  |

To test for within-species variability, most species were represented by multiple origins, further called accessions. The number of accessions per species was driven by seed availability and varied strongly between species. More than half of the species were represented by at least four origins, while for six species we managed to obtain only one accession (Table 1). In total, the experiment included 188 accessions.

The experiment required large amounts of seeds. Seeds of 14 accessions were obtained by direct collection in nature or by collecting at least 50 young plants in the wild and growing them in a cold greenhouse to produce seeds. However, most seeds come from on-farm propagation for restoration purposes. We are aware that on-farm propagation may cause unintended selection, which could alter seed traits (Conrady et al., 2023; Ensslin et al., 2018). In the seeds we used, this effect will be rather minor because most producers are relatively new in this business and the propagation did not take place for many generations (*personal communication*).

Using farm-propagated seeds further has the advantage that the maternal plants grew under optimal conditions and the seeds were harvested when fully ripe. This reduces the variability in seed quality, a characteristic that has a massive effect on storage behaviour (Rahman & Ellis, 2019; Sano et al., 2016; Sinniah et al., 1998).

### Artificial ageing

To artificially age the seeds, we exposed them to 45° C and 60% relative humidity in a climate-controlled cabinet (Rumed ®, Rubarth Apparate GmbH, Laatzen, Germany). These conditions enhance similar chemical processes that are typical of natural ageing during long-term storage and accelerate seed deterioration (Merritt et al., 2014; Probert et al., 2009).

We exposed the seeds to artificial ageing conditions for 0 (control group, fresh seeds), 1, 5, 9, 15, 20, 30, 40, 57 and 72 days (adjusted after Probert et al. (2009)). We put 3 replicates of 30 seeds of every accession and ageing duration into open 1.5 mL tubes in the ageing condition. The position of each accession in the ageing condition was randomised within the ageing duration. Before the ageing treatment, we manually scarified Fabaceae seeds with sandpaper to allow for moisture to be imbibed (Baskin, 2003). In total, our experiment contained 188 accessions, each with 10 ageing treatments and 3 replicates, that is 5,640 replicates containing 30 seeds each, that is 169,200 seeds.

### Germination test

For germination tests, we used 6-well plates with a well-diameter of 39 mm (VWR International GmbH, Darmstadt, Germany, catalogue number: 734-2323), because they use space in the climate cabinet more efficiently than single petri dishes. For species with large seeds (*Bupleurum rotundifolium*, *Centaurea cyanus*, *Bromus hordeaceus*, *Arrhenatherum elatius*, *Lathyrus nissolia*, *Hordeum murinum*, *Hypochaeris radicata*, *Agrostemma githago*, *Poa annua*) we used petri dishes of 6 cm diameter to avoid overcrowding. We wrapped Petri dishes with seeds of *Bromus hordeaceus* in aluminium foil since light slightly inhibits germination (Andersson et al., 2002). After the intended ageing duration, we removed the seeds from the ageing conditions and transferred each replicate into one well or a petri dish laid out with two filter papers in a fully randomised design. We watered the seeds with a 0.025% Gibberellic acid solution to break dormancy and closed the plates with transparent Parafilm® to avoid evaporation. Germination took place in a climate-controlled cabinet (Rumed®, Rubarth Apparate GmbH, Laatzen, Germany) with a 14 h daily photoperiod at 20° C and 10 h darkness at 10° C.

We checked for germination every seven days and added distilled water if necessary. When a radicle was longer than 2 mm, we considered a seed viable and removed it from the dish. For each replicate with viable seeds, we finished the trial when no more seeds germinated for at least four weeks. If in a replicate no seeds germinated at all for ten weeks, we considered the seeds not viable. If samples were affected by mould, we added 1 ml of a 0.0835% Difenoconazol solution.

### Seed traits and climatic data

For each accession, we determined seed weight by weighing 2 replicates of 50 seeds and calculated the weight of a thousand seeds (further called seed weight for simplicity). We also obtained the mean annual temperature and precipitation for each accession location from WorldClim2 using the R package *raster* (Fick & Hijmans, 2017). In wild-collected seeds, we used the exact location of the seed collection. For farm-propagated seeds, we used the location of the farm, as the initial seeds used to establish the seed orchard were typically collected in the same region where the seed producer is located (Bucharova et al., 2019, Figure 1). The oil and protein content of the seeds was obtained from the Seed Information Database on species level (Society for Ecological Restoration et al., 2023).

**Figure 1:**
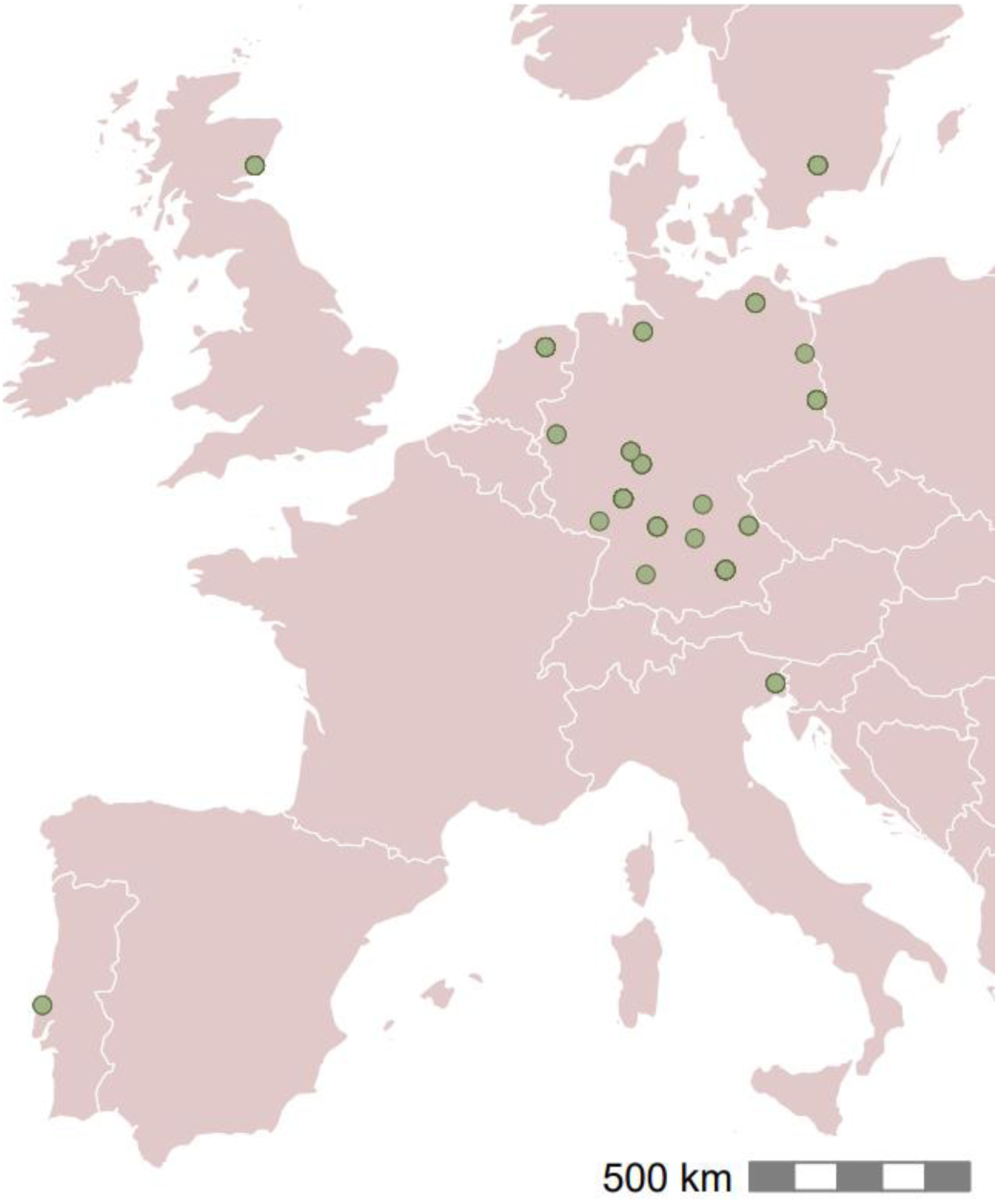
Origins of the 135 seed accessions used in the study and included in the analyses. Each point represents the location of a seed producer as a proxy for the origins of the individual seed accessions and thus, is the origin of multiple species. In three cases, the points represent the location of own seed collection.

### Statistical analyses

Before data analysis, we calculated the initial germination rate of each of the 188 seed accessions. In some species, initial germination rates were higher in short-duration ageing treatments (one or five days) than in fresh seeds. This was likely because short exposure to ageing treatment can break dormancy (Hay et al., 2021; Mira et al., 2019). We thus defined “initial germination” for each accession as the highest average germination rate of the 0, 1 or 5 days in the ageing treatment. We removed accessions with initial germination below 50% and those particularly affected by mould (n = 39).

To analyse seed survival, we used generalized linear models (GLMs) with a quasibinomial error distribution (to account for overdispersion) and a probit link function. The probit transformation converts percentage responses onto a linear scale with a normal error distribution. We modelled the proportion of germinated seeds per replicate as a function of the number of days in ageing (Wolkis et al., 2025). The slope of the probit function directly describes the rate of seed viability loss, and back transformation of the slope (−1/slope) yields σ, the number of days needed for viability to drop by one probit (Hay et al., 2022; Klepka et al., 2025). We are aware that most comparative studies of seed longevity in storage measured seed longevity as *p*_50_, the number of days needed for viability to decrease to 50%, yet this parameter is heavily dependent on the initial seed viability (see Klepka et al. (2025) for details). While the initial viability can be affected by the environmental effects, e.g. during seed storage before the experiment, σ is assumed to be a genetic trait that describes a species, population or genotype, and is thus the actual trait of interest (Hay et al., 2022). We thus decided to use σ as a measure of seed longevity, because it is independent of the initial seed viability of given accessions and directly describes the slope of seed viability loss (Hay et al., 2022; Klepka et al., 2025).

## Within-species variability

We obtained the seed longevity estimate (σ) for each accession. To do this, we related seed germination in each replicate to the time in ageing conditions in a separate probit model for each accession. If the model failed to give significant estimates for the slope, or if σ was higher than the maximum ageing duration in our experiment (72 days), we excluded the accession from further analyses (n = 14), since we assumed that in these cases the data cover only a very short portion of the survival curve and the estimated parameters are therefore unprecise. These models yielded separate σ-values for each accession.

We then tested whether the seed longevity (σ) differs between accessions within species. This was possible for 26 species for which we had multiple accessions that passed the initial check (initial germination >50% and significant accession-specific σ). For this, we related the seed germination in each replicate to the accession identity, the ageing duration and their interaction as explanatory variables in a separate probit model for each species. A significant interaction between accession identity and ageing duration indicates significantly different slopes between accessions within species and therefore different seed longevities (σ). We further tested whether the probability of detecting significant differences in σ between accessions within species depended on the number of accessions representing each species in our data set. To do this, we related the presence of significant difference between accessions in each species (binomial yes/no) to the number of accessions per species in a generalised linear model with binomial error distribution.

To understand drivers of within-species variability in seed longevity in storage, we related accession-specific longevity (σ) as the response variable to the accession-specific seed weight, mean annual temperature and precipitation at the accession’s origin as explanatory variables in a multiple linear mixed model. We included the species’ identity as a random factor. For this analysis, we scaled the seed weight within each species to standardise the variable across species, and included only species that were represented by at least 2 accessions.

We quantified the proportion of within-species variance explained by the fixed effects using marginal and conditional R^2^ estimates from the MuMIn package. This allowed us to calculate the contribution of the within-species predictors relative to the variance remaining after accounting for species identity.

## Between-species variability

We obtained the seed longevity estimate (σ) for each species, common across accessions, regardless whether we detected significant differences among accessions in the previous step. For this, we related seed germination in each replicate (and from all accession of given species) to time in ageing in a probit model for each species. For species with more than one accession, we added accession identity as an explanatory variable to allow for different intercepts (initial viability) in individual accessions. These models yielded one σ for each species. To understand drivers of between-species variability in seed longevity in storage, we related the log-transformed species-specific σ as the response variable to the species’ mean seed weight and plant family as explanatory variables in a multiple linear regression model. As protein and oil contents were only available for 15 study species, we evaluated their effects in a separate multiple linear regression model.

All data analyses were performed using R version 4.4.2 (2024-10-31 ucrt) (R Development Core Team, 2024). We log-transformed σ as the response variable to ensure that model assumptions were met. For all models, we verified that model assumptions were met by graphically evaluating the residuals (Zuur et al., 2010).

## RESULTS

The estimated seed longevity (σ) significantly varied among accessions within 58% of the species for which we had multiple accessions, specifically in 15 out of 26 species (Table S1). The probability of detecting significant differences among accessions within species increased with the number of included accessions (R^2^=0.20, p=0.016, Figure S1). Within-species variation was the highest in *Prunella vulgaris*, where σ varied more than 4-fold (Figure 2), specifically between 12.66 and 53.14 days. This within-species variability was hard to predict, as accession-specific predictors explained only 14.4% of the variability among accessions, after correcting for species identity. Specifically, seed longevity decreased with seed weight and increased with the mean annual temperature at the accession origin, but was unrelated to the mean annual precipitation (Table 2, Figure 3).

**Figure 2:**
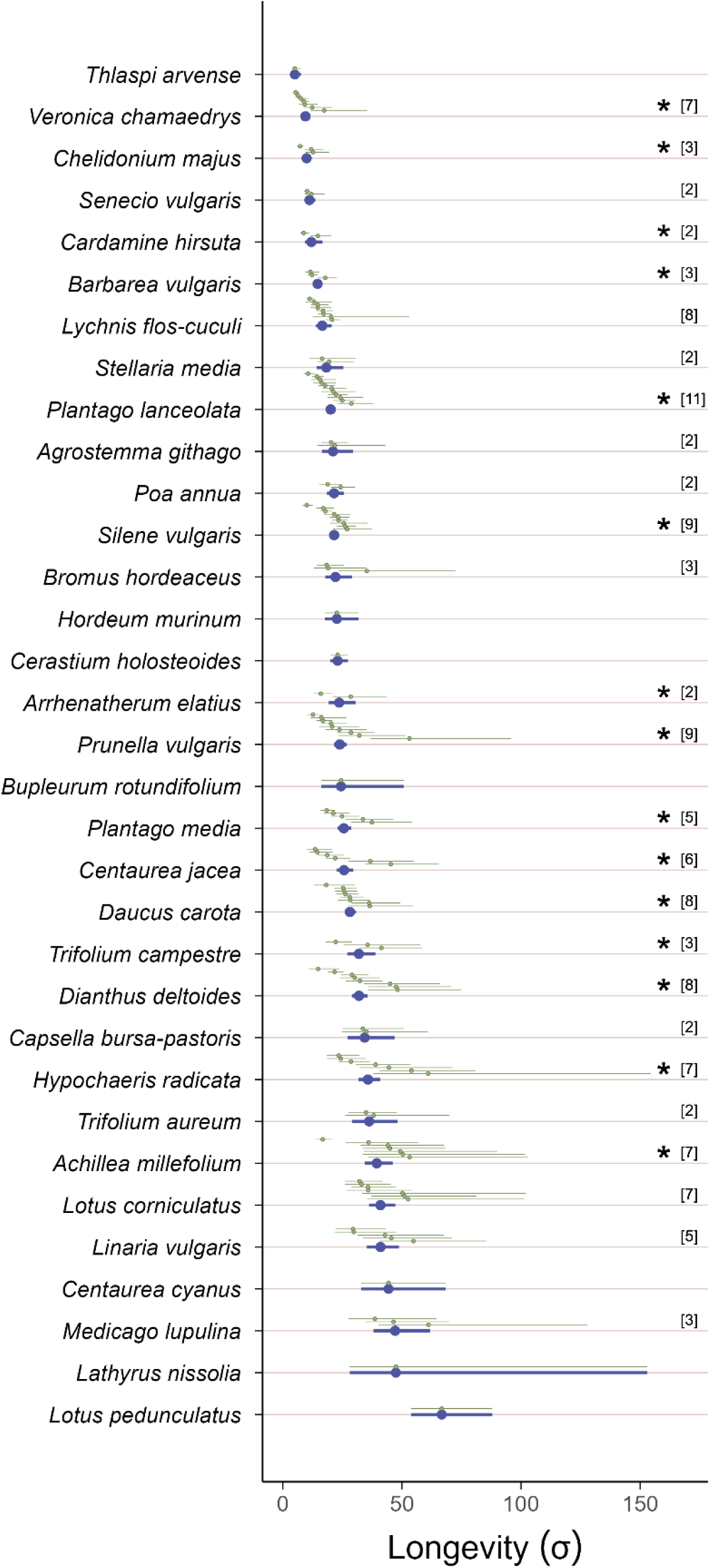
Estimated seed longevity (σ) and 95% confidence intervals for 33 species and 135 accessions. Blue points and bars represent species-level estimates; green points and bars show accession-level estimates. Asterisks mark significant differences between accessions within species. Numbers in brackets indicate the number of accessions for species represented by multiple accessions.

**Figure 3:**
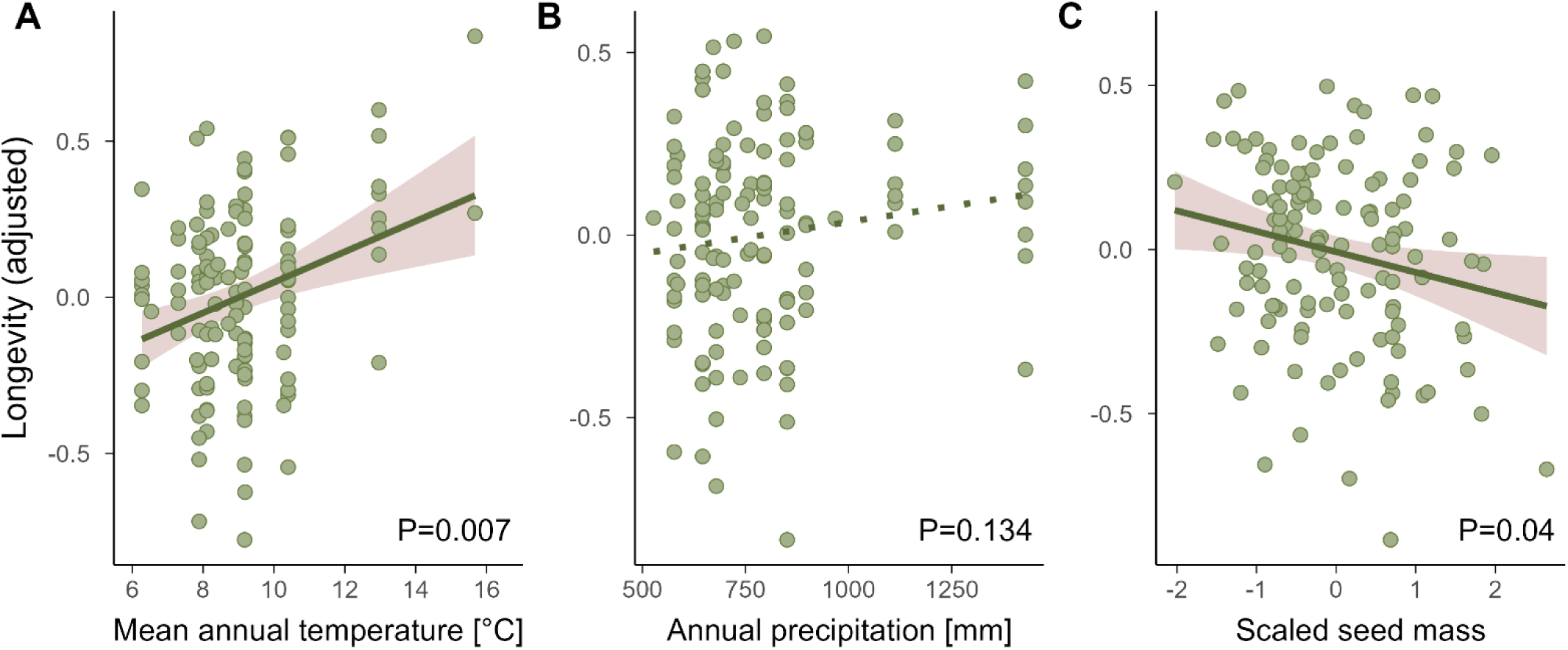
Relationship between seed longevity (σ) of each accession, accession-specific seed traits, and environmental variables. Each point represents a seed accession. Seed longevity was adjusted for species identity. (A) Mean annual temperature, (B) annual precipitation at the origins of the accessions, and (C) accession-specific seed mass (scaled); For full model results, see Table 2.

**Table 2:** Intraspecific variability in seed longevity in storage, results of the linear mixed model testing the effect of accession-specific seed weight, mean annual temperature, and annual precipitation on accession-specific seed longevity in storage with species identity as random factor. Analysis of variance with error type III. Significant effects (P < 0.05) are highlighted in bold.

|  | df | Chi Sq | p-value | margR<sup>2</sup> | condR<sup>2</sup> |
| --- | --- | --- | --- | --- | --- |
|  |  |  |  | 0.05 | 0.72 |
| Accession-specific seed weight | 1 | 4.23 | **0.040** |  |  |
| Mean annual temperature | 1 | 7.30 | **0.007** |  |  |
| Annual precipitation | 1 | 2.25 | 0.134 |  |  |

The estimated seed longevity also varied among species (Figure 2), and species s identity explained 66% of the total variability among accessions. The average seed longevity was shortest for *Thlaspi arvense* (σ = 5.04 days) and longest for *Lotus pedunculatus* (σ = 66.70 days). The species seed longevity was unrelated to the species-specific seed weight (Table 3A, Figure 4A) or the seed oil and protein content (Table 3B, Figure 4B, C), but we found significant differences between plant families (Table 3A, Figure 4D).

**Figure 4:**
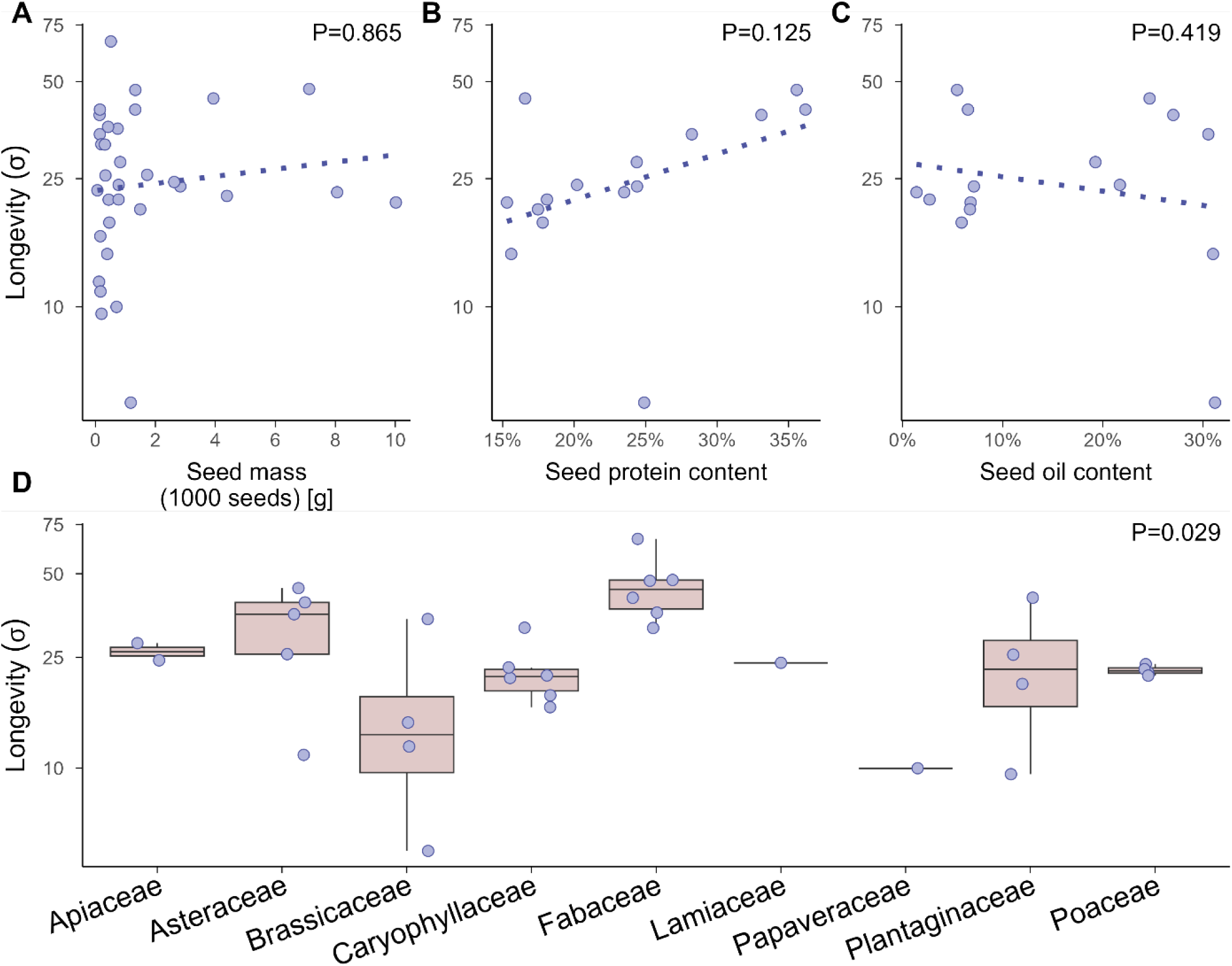
Relationship between species seed longevity (σ) and species-specific seed traits. Each point represents one species. (A) Seed mass, average across accessions of the same species; species-specific (B) protein and (C) oil content in the seeds; and (D) plant families. The dashed lines indicate non-significant relationships. For full model results, see Table 3.

**Table 3:** Interspecific variability in seed longevity in storage, results of the linear model testing the effect of (A) the family, the species-specific seed weight, and (B) the protein and oil content on species-specific seed longevity. Analysis of variance with error type III. Significant effects (p < 0.05) are highlighted in bold.

|  | df | Resid.df | *F* | *p* | R<sup>2</sup> |
| --- | --- | --- | --- | --- | --- |
| A |  | 23 |  |  | 0.49 |
| Family | 8 |  | 2.72 | **0.029** |  |
| Species-specific seed weight | 1 |  | 0.03 | 0.865 |  |
| B |  | 12 |  |  | 0.22 |
| Protein content | 1 |  | 0.77 | 0.125 |  |
| Oil content | 1 |  | 0.70 | 0.419 |  |

## DISCUSSION

Effective conservation of seeds in ex-situ seed banks requires an understanding of seed storage behaviour not only for individual species but also for accessions within the very same species. We found, for multiple species common in European grasslands, that accessions of the same species commonly differ in seed longevity in storage. This intraspecific variability was partially explained by seed weight and mean annual temperature at the accession origin, but the predictive power of these variables was rather low (2.1% explained variability across all species). On the other hand, species identity was a strong predictor of seed longevity, explaining 66% of the variability among accessions. Our results thus suggest that seed longevity can be reliably predicted at the species level, but predicting accession-specific seed longevity in storage remains challenging.

### Intraspecific variability in seed longevity

Seed longevity in storage varied considerably within-species, with accession-specific seed longevity differing significantly in 15 out of 26 species with multiple accessions. The probability of detecting significant differences among accessions within a species increased with the number of accessions included in the study. This suggests that with more accessions available, significant differences would likely be detected in even more species. Consequently, within-species differences in seed longevity in storage are likely very common.

Within-species, seed longevity was partially explained by the mean annual temperature at the collection site, a pattern that was reported previously (Mondoni et al., 2011). Other studies attributed such a positive correlation between temperature and seed longevity mainly to the species identity because they had species represented by single accessions (Merritt et al., 2014; Probert et al., 2009). We did not find any effect of the annual precipitation on seed longevity, which contrasts with previous studies, both within and across species (Merritt et al., 2014; Probert et al., 2009; White et al., 2023). This is likely because the mean annual precipitation was correlated with mean annual temperature (r=0.34), and as we used multivariate models, we were not able to tease apart the effect of both climate variables.

We found a negative relationship between the accession-specific seed weight and seed longevity, with lighter seeds surviving significantly longer. Previous studies primarily tested the relationship between seed weight and longevity across species, yielding mixed results (Davies et al., 2020; Merritt et al., 2014; Pritchard & Dickie, 2003; Probert et al., 2009; Satyanti et al., 2018). Similarly, research on variability within individual species has also produced inconsistent findings (Franks et al., 2018; Genna et al., 2020; Guzzon et al., 2018, 2021; Mira et al., 2019; Schutte et al., 2008). These inconsistent study findings underscore that seed mass remains an unreliable predictor and highlight the complexity of seed longevity and its drivers. Nevertheless, our results provide new evidence suggesting that, for a variety of wild grassland species, lighter seeds may have a slight but consistent advantage over heavier seeds of the same species in long-term storage survival.

The intraspecific variability in seed longevity may be driven by two mutually non-exclusive mechanisms – phenotypic plasticity or genetic differentiation. For example, Mondoni et al. (2011) detected that seeds collected in populations in warmer environments survived longer in artificial ageing than conspecific seeds collected in cooler environments. A large proportion of this effect was caused by phenotypic plasticity, but the pattern was still significant, although weaker, in the second generation grown in a common environment, which indicates that the effect is partially heritable and thus genetic (Mondoni et al., 2014). Indeed, genes associated with seed longevity in storage were identified in some crops and model species (Bizouerne et al., 2023; Nagel et al., 2015; Raquid et al., 2021; Renard et al., 2020). Our study worked with seeds that were collected in natural sites or produced in the region of origin; we thus cannot discriminate whether the observed effects are plastic or genetic.

After accounting for species identity, within-species predictors explained 14.4% of the remaining variability among accessions. Still, species identity was the strongest predictor of seed longevity and explained 66.9%. Our findings align with those of Kochanek et al. (2009). On a continental scale, they collected data on seed longevity in seven species represented by 2-8 populations, with populations within species largely aggregated in a specific biome. Here, the species identity explained 69.1% (adjusted R^2^) of the variability among accessions, leaving 30% to variation among accessions within species (own analysis of the data presented in Kochanek et al. (2009)). In contrast, a study by Mondoni et al. found that the intraspecific variability in seed longevity was much larger than the interspecific one. On a regional scale, they examined the seed longevity of six ecologically similar species that were each represented by two populations in contrasting environments, specifically high and low altitudes. Here, species identity explained only 6% of variability, leaving an astonishing 94% variation within species (own analysis of the data presented in Mondoni et al. (2011)). In our study, we detected that species explained about two-thirds of the variability in seed longevity among accessions, yet this number might be underestimated because we excluded the species with the most long-lived seeds for methodical reasons (see methods).

The comparison of our results with the findings of previous studies is complicated by different measures of seed longevity. We used σ, the rate of seed viability loss, which is independent of the initial seed viability and thus suitable for comparative studies across accessions and species. The vast majority of previous studies used *p*_50_, the time till the seed viability decreases to half (e.g. Mondoni et al., 2011; Probert et al., 2009; Satyanti et al., 2018), a measure that is partially determined by the rate of seed viability loss (σ), but is heavily dependent on initial seed viability at the start of the experiment (Klepka et al., 2025). For example, using seeds with an initial viability ranging from 85 to 99.9%, but with the same rate of seed viability loss (σ), results in a threefold variation in *p*_50_. The *p*_50_ estimate thus does not appropriately represent inherent differences between genotypes or populations (Hay et al., 2019). Direct comparison between our study and many previous studies is thus complex, unless raw viability data are available to calculate σ.

### Interspecific variability in seed longevity

Across species, the only variable that significantly affected seed longevity was the plant family, with the observed differences between families being roughly in line with previous studies (Merritt et al., 2014; Mondoni et al., 2011; Probert et al., 2009; Walters et al., 2005b): Fabaceae seeds were relatively long-lived, probably due to their physical dormancy (Merritt et al., 2014), there was a large variability within Asteraceae (Merritt et al., 2014; Walters et al., 2005a), and three of the six shortest-lived species in our study belonged to the Brassicaceae family (*Thlaspi arvense*, *Cardamine hirsuta*, *Barbarea vulgaris*).

Neither seed weight nor chemical composition influenced seed longevity in our artificial ageing experiment. In general, species-specific seed weight seems to be an unreliable predictor of seed longevity in storage, as the effects reported in previous literature strongly vary from negative, over no effect to, rarely, positive (Davies et al., 2020; Merritt et al., 2014; Pritchard & Dickie, 2003; Probert et al., 2009; Satyanti et al., 2018). Similarly, the chemical composition of the seeds did not predict seed longevity, which corresponds to previous empirical studies of wild species (Kochanek et al., 2009; Walters et al., 2005b). The expected effect of the chemical composition on seed longevity in storage, especially oil content, is based on theoretical expectations and comparisons among oil crops, cereals, and legumes (Narayana Murthy & Sun, 2000; Pritchard & Dickie, 2003 Nagel et al., 2015), but the empirical evidence provided by others and us shows that it is likely not true for wild species.

## CONCLUSIONS

Our study demonstrates that in the majority of species, there is substantial within-species variation in the rate of seed viability loss in storage. Conservation seed banks sometimes design viability monitoring of individual accessions based on species identity and initial seed viability only, assuming that accessions of the same species lose viability at the same rate when stored under the same conditions (Hay & Whitehouse, 2017). Our data show that this assumption is not valid and support the existing concerns that this method might be suboptimal.

## ACKNOWLEDGEMENTS

We thank Johannes Kasper, Lena Lerbs, Christina Mengel, Sarah Paschen, Nadine Pluquette, Annika Sommerfeld, and Helene Villhauer for technical assistance. This project was funded by a scholarship of Marburg University Research Academy (MARA) to LK.

## AUTHOR CONTRIBUTIONS

Lea Klepka: Investigation, Formal analysis, Data Curation, Writing – original Draft, Visualization, Funding Acquisition. Sacha Liepelt: Investigation, Writing – Review&Editing, Anna Bucharova: Conceptualization, Methodology, Resources, Writing – Review & Editing, Supervision

## DATA AVAILABILITY STATEMENT

The data supporting the findings of this study are not publicly available at the time of submission but will be deposited in an appropriate public repository upon acceptance of the manuscript. Data will be made available to editors and reviewers upon reasonable request.

---

**Figure S1:**
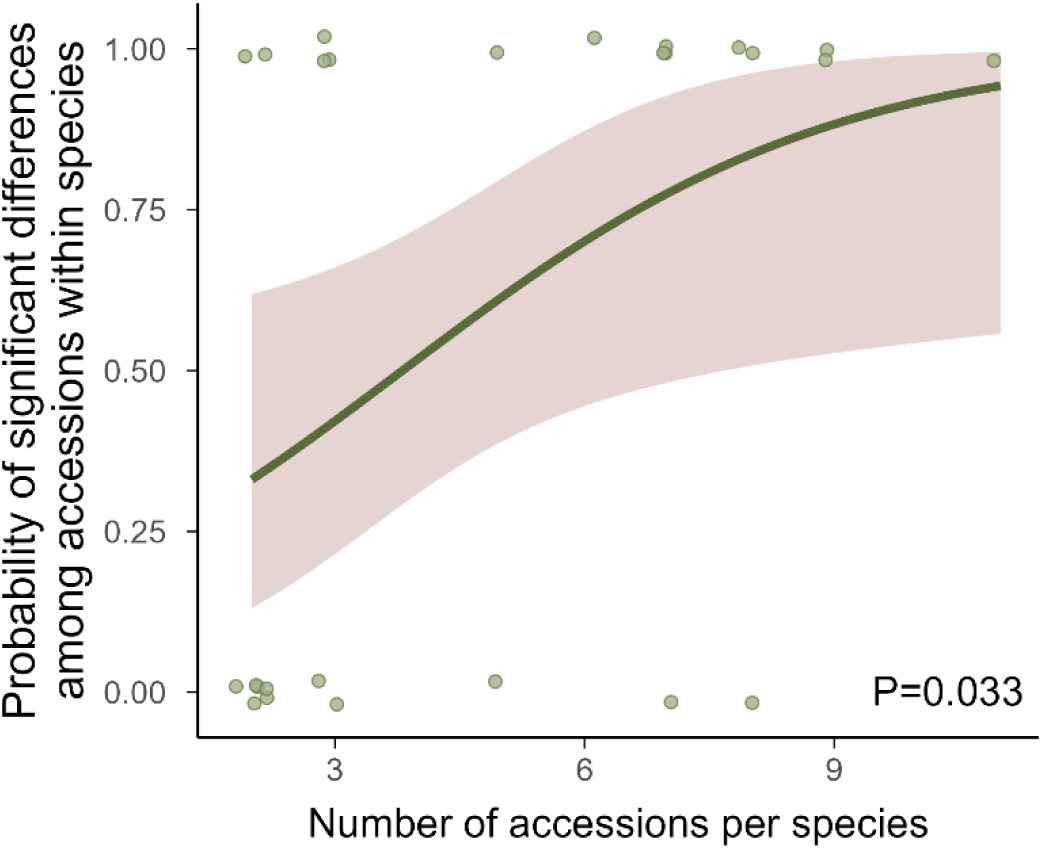
Relationship between the number of accessions per species and the probability of detecting significant differences among accessions within species. The line shows a logistic regression with the 95% confidence interval.

**Table S1:** Species-specific seed survival models. Bold type indicates a significant effect of the parameter. A significant interaction between the accession identity and the ageing duration means significantly different seed longevities between accessions within species.

| Species | Number of Accessions | Ageing Duration |  |  | Accession Identity |  |  | Accession × Ageing |  |  |
| --- | --- | --- | --- | --- | --- | --- | --- | --- | --- | --- |
|  |  | Df | F-value | p-value | Df | F-value | p-value | Df | F-value | p-value |
| *Achillea millefolium* | 7 | 1 | 20.05 | **<0.001** | 6 | 1.24 | 0.290 | 6 | 5.96 | **<0.001** |
| *Agrostemma githago* | 2 | 1 | 37.33 | **<0.001** | 1 | 0.24 | 0.623 | 1 | 0.07 | 0.788 |
| *Arrhenatherum elatius* | 2 | 1 | 66.81 | **<0.001** | 1 | 1.10 | 0.298 | 1 | 6.41 | **0.014** |
| *Barbarea vulgaris* | 3 | 1 | 124.86 | **<0.001** | 2 | 56.90 | **<0.001** | 2 | 5.52 | **0.006** |
| *Bromus hordeaceus* | 3 | 1 | 16.95 | **<0.001** | 2 | 11.07 | **<0.001** | 2 | 2.49 | 0.089 |
| *Capsella bursa-pastoris* | 2 | 1 | 30.59 | **<0.001** | 1 | 1.98 | 0.165 | 1 | 0.03 | 0.871 |
| *Cardamine hirsuta* | 2 | 1 | 194.27 | **<0.001** | 1 | 4.24 | **0.044** | 1 | 8.65 | **0.005** |
| *Centaurea jacea* | 6 | 1 | 50.24 | **<0.001** | 5 | 7.59 | **<0.001** | 5 | 9.85 | **<0.001** |
| *Chelidonium majus* | 3 | 1 | 101.61 | **<0.001** | 2 | 19.83 | **<0.001** | 2 | 5.23 | **0.007** |
| *Daucus carota* | 8 | 1 | 84.95 | **<0.001** | 7 | 11.50 | **<0.001** | 7 | 2.50 | **0.017** |
| *Dianthus deltoides* | 8 | 1 | 99.55 | **<0.001** | 7 | 5.94 | **<0.001** | 7 | 8.28 | **<0.001** |
| *Hypochaeris radicata* | 7 | 1 | 80.70 | **<0.001** | 6 | 10.98 | **<0.001** | 6 | 5.19 | **<0.001** |
| *Linaria vulgaris* | 5 | 1 | 23.02 | **<0.001** | 4 | 11.62 | **<0.001** | 4 | 2.09 | 0.086 |
| *Lotus corniculatus* | 7 | 1 | 26.03 | **<0.001** | 6 | 4.06 | **0.001** | 6 | 1.58 | 0.155 |
| *Lychnis flos-cuculi* | 8 | 1 | 82.17 | **<0.001** | 7 | 18.94 | **<0.001** | 7 | 1.98 | 0.059 |
| *Medicago lupulina* | 3 | 1 | 23.12 | **<0.001** | 2 | 9.84 | **<0.001** | 2 | 1.21 | 0.305 |
| *Plantago lanceolata* | 11 | 1 | 123.03 | **<0.001** | 10 | 12.22 | **<0.001** | 10 | 5.14 | **<0.001** |
| *Plantago media* | 5 | 1 | 56.65 | **<0.001** | 4 | 22.08 | **<0.001** | 4 | 5.59 | **<0.001** |
| *Poa annua* | 2 | 1 | 94.97 | **<0.001** | 1 | 0.02 | 0.877 | 1 | 2.47 | 0.122 |
| *Prunella vulgaris* | 9 | 1 | 37.96 | **<0.001** | 8 | 4.85 | **<0.001** | 8 | 6.50 | **<0.001** |
| *Senecio vulgaris* | 2 | 1 | 149.25 | **<0.001** | 1 | 2.91 | 0.093 | 1 | 0.81 | 0.372 |
| *Silene vulgaris* | 9 | 1 | 139.81 | **<0.001** | 8 | 12.98 | **<0.001** | 8 | 7.27 | **<0.001** |
| *Stellaria media* | 2 | 1 | 46.77 | **<0.001** | 1 | 1.25 | **0.268** | 1 | 0.31 | 0.579 |
| *Trifolium aureum* | 2 | 1 | 29.80 | **<0.001** | 1 | 5.75 | **0.020** | 1 | 0.12 | 0.732 |
| *Trifolium campestre* | 3 | 1 | 87.20 | **<0.001** | 2 | 2.68 | 0.074 | 2 | 4.83 | **0.010** |
| *Veronica chamaedrys* | 7 | 1 | 128.71 | **<0.001** | 6 | 10.05 | **<0.001** | 6 | 5.46 | **<0.001** |

## REFERENCES

Andersson, L., Milberg, P., Schütz, W., & Steinmetz, O. (2002). Germination characteristics and emergence time of annual Bromus species of differing weediness in Sweden. Weed Research, 42(2), 135–147. 10.1046/j.1365-3180.2002.00269.x

Baskin, C. C. (2003). Breaking physical dormancy in seeds – focussing on the lens. New Phytologist, 158(2), 229–232. 10.1046/j.1469-8137.2003.00751.x

Bizouerne, E., Ly Vu, J., Ly Vu, B., Diouf, I., Bitton, F., Causse, M., Verdier, J., Buitink, J., & Leprince, O. (2023). Genetic Variability in Seed Longevity and Germination Traits in a Tomato MAGIC Population in Contrasting Environments. *Plants (Basel*, Switzerland), 12(20), 3632. 10.3390/plants12203632

Bucharova, A., Bossdorf, O., Hölzel, N., Kollmann, J., Prasse, R., & Durka, W. (2019). Mix and match: Regional admixture provenancing strikes a balance among different seed-sourcing strategies for ecological restoration. Conservation Genetics, 20(1), 7–17. 10.1007/s10592-018-1067-6

Cochrane, J. A., Crawford, A. D., Monks, L. T., Cochrane, J. A., Crawford, A. D., & Monks, L. T. (2007). The significance of ex situ seed conservation to reintroduction of threatened plants. Australian Journal of Botany, 55(3), 356–361. 10.1071/BT06173

Cohen, J. I., Williams, J. T., Plucknett, D. L., & Shands, H. (1991). Ex Situ Conservation of Plant Genetic Resources: Global Development and Environmental Concerns. Science, 253(5022), 866–872. 10.1126/science.253.5022.866

Conrady, M., Lampei, C., Bossdorf, O., Hölzel, N., Michalski, S., Durka, W., & Bucharova, A. (2023). Plants cultivated for ecosystem restoration can evolve toward a domestication syndrome. Proceedings of the National Academy of Sciences, 120(20), e2219664120. 10.1073/pnas.2219664120

Davies, R. M., Hudson, A. R., Dickie, J. B., Cook, C., O’Hara, T., & Trivedi, C. (2020). Exploring seed longevity of UK native trees: Implications for *ex situ* conservation. Seed Science Research, 30(2), 101–111. 10.1017/S0960258520000215

Delouche, J., & Baskin, C. (2021). Accelerated Aging Techniques for Predicting the Relative Storability of Seed Lots. All Articles. https://scholarsjunction.msstate.edu/seedtechpapers/10

ENSCONET. (2009, June 16). ENSCONET Curation Protocols & Recommendations. http://ensconet.maich.gr/PDF/Curation_protocol_English.pdf

Ensslin, A., Van de Vyver, A., Vanderborght, T., & Godefroid, S. (2018). Ex situ cultivation entails high risk of seed dormancy loss on short-lived wild plant species. Journal of Applied Ecology, 55(3), 1145–1154. 10.1111/1365-2664.13057

Etterson, J. R., Franks, S. J., Mazer, S. J., Shaw, R. G., Gorden, N. L. S., Schneider, H. E., Weber, J. J., Winkler, K. J., & Weis, A. E. (2016). Project Baseline: An unprecedented resource to study plant evolution across space and time. American Journal of Botany, 103(1), 164–173. 10.3732/ajb.1500313

Fick, S. E., & Hijmans, R. J. (2017). WorldClim 2: New 1-km spatial resolution climate surfaces for global land areas. International Journal of Climatology, 37(12), 4302–4315. 10.1002/joc.5086

Franks, S. J., Hamann, E., & Weis, A. E. (2018). Using the resurrection approach to understand contemporary evolution in changing environments. Evolutionary Applications, 11(1), 17–28. 10.1111/eva.12528

Franks, S. J., Sekor, M. R., Davey, S., & Weis, A. E. (2019). Artificial seed aging reveals the invisible fraction: Implications for evolution experiments using the resurrection approach. Evolutionary Ecology, 33(6), 811–824. 10.1007/s10682-019-10007-2

Genna, N. G., Walters, C., & Pérez, H. E. (2020). Viability and vigour loss during storage of Rudbeckia mollis seeds having different mass: An intra-specific perspective. Seed Science Research, 30(2), 122–132. 10.1017/S0960258520000161

Guzzon, F., Gianella, M., Velazquez Juarez, J. A., Sanchez Cano, C., & Costich, D. E. (2021). Seed longevity of maize conserved under germplasm bank conditions for up to 60 years. Annals of Botany, 127(6), 775–785. 10.1093/aob/mcab009

Guzzon, F., Orsenigo, S., Gianella, M., Müller, J. V., Vagge, I., Rossi, G., & Mondoni, A. (2018). Seed heteromorphy influences seed longevity in Aegilops. Seed Science Research, 28(4), 277–285. 10.1017/S096025851800034X

Hay, F. R., Davies, R. M., Dickie, J. B., Merritt, D. J., & Wolkis, D. M. (2022). More on seed longevity phenotyping. Seed Science Research, 32(3), 144–149. 10.1017/S0960258522000034

Hay, F. R., & Probert, R. J. (2013). Advances in seed conservation of wild plant species: A review of recent research. Conservation Physiology, 1(1), cot030–cot030. 10.1093/conphys/cot030

Hay, F. R., Valdez, R., Lee, J.-S., & StaCruz, P. C. (2019). Seed longevity phenotyping: Recommendations on research methodology. Journal of Experimental Botany, 70(2), 425–434. 10.1093/jxb/ery358

Hay, F. R., & Whitehouse, K. J. (2017). Rethinking the approach to viability monitoring in seed genebanks. Conservation Physiology, 5(1), cox009. 10.1093/conphys/cox009

Hay, F. R., Whitehouse, K. J., Ellis, R. H., Sackville Hamilton, N. R., Lusty, C., Ndjiondjop, M. N., Tia, D., Wenzl, P., Santos, L. G., Yazbek, M., Azevedo, V. C. R., Peerzada, O. H., Abberton, M., Oyatomi, O., de Guzman, F., Capilit, G., Muchugi, A., & Kinyanjui, Z. (2021). CGIAR genebank viability data reveal inconsistencies in seed collection management. Global Food Security, 30, 100557. 10.1016/j.gfs.2021.100557

Klepka, L., Carta, A., Hay, F. R., & Bucharova, A. (2025). Limitations of p50 as a measure of seed longevity and the way forward (p. 2025.06.11.659081). bioRxiv. 10.1101/2025.06.11.659081

Kochanek, J., Buckley, Y. M., Probert, R. J., Adkins, S. W., & Steadman, K. J. (2010). Pre-zygotic parental environment modulates seed longevity. Austral Ecology, 35(7), 837–848. 10.1111/j.1442-9993.2010.02118.x

Kochanek, J., Steadman, K. J., Probert, R. J., & Adkins, S. W. (2009). Variation in seed longevity among different populations, species and genera found in collections from wild Australian plants. Australian Journal of Botany, 57(2), 123–131. 10.1071/BT09023

Lee, J.-S., Velasco-Punzalan, M., Pacleb, M., Valdez, R., Kretzschmar, T., McNally, K. L., Ismail, A. M., Cruz, P. C. S., Sackville Hamilton, N. R., & Hay, F. R. (2019). Variation in seed longevity among diverse Indica rice varieties. Annals of Botany, 124(3), 447–460. 10.1093/aob/mcz093

Merritt, D. J., & Dixon, K. W. (2011). Restoration Seed Banks—A Matter of Scale. Science, 332(6028), 424–425. 10.1126/science.1203083

Merritt, D. J., Martyn, A. J., Ainsley, P., Young, R. E., Seed, L. U., Thorpe, M., Hay, F. R., Commander, L. E., Shackelford, N., Offord, C. A., Dixon, K. W., & Probert, R. J. (2014). A continental-scale study of seed lifespan in experimental storage examining seed, plant, and environmental traits associated with longevity. Biodiversity and Conservation, 23(5), 1081–1104. 10.1007/s10531-014-0641-6

Millennium Seed Bank. (2022). Assessing a Population for Seed Collection. https://brahmsonline.kew.org/Content/Projects/msbp/resources/Training/02-Assessing-population.pdf

Mira, S., Veiga-Barbosa, L., & Pérez-García, F. (2019). Seed dormancy and longevity variability of *Hirschfeldia incana* L. during storage. Seed Science Research, 29(2), 97–103. 10.1017/S0960258519000072

Mondoni, A., Orsenigo, S., Donà, M., Balestrazzi, A., Probert, R. J., Hay, F. R., Petraglia, A., & Abeli, T. (2014). Environmentally induced transgenerational changes in seed longevity: Maternal and genetic influence. Annals of Botany, 113(7), 1257–1263. 10.1093/aob/mcu046

Mondoni, A., Probert, R. J., Rossi, G., Vegini, E., & Hay, F. R. (2011). Seeds of alpine plants are short lived: Implications for long-term conservation. Annals of Botany, 107(1), 171–179. 10.1093/aob/mcq222

Nagel, M., Kranner, I., Neumann, K., Rolletschek, H., Seal, C. E., Colville, L., Fernández-Marín, B., & Börner, A. (2015). Genome-wide association mapping and biochemical markers reveal that seed ageing and longevity are intricately affected by genetic background and developmental and environmental conditions in barley. Plant, Cell & Environment, 38(6), 1011–1022. 10.1111/pce.12474

Narayana Murthy, U. M., & Sun, W. Q. (2000). Protein modification by Amadori and Maillard reactions during seed storage: Roles of sugar hydrolysis and lipid peroxidation. Journal of Experimental Botany, 51(348), 1221–1228. 10.1093/jexbot/51.348.1221

Niedzielski, M., Walters, C., Luczak, W., Hill, L. M., Wheeler, L. J., & Puchalski, J. (2009). Assessment of variation in seed longevity within rye, wheat and the intergeneric hybrid triticale. Seed Science Research, 19(4), 213–224. 10.1017/S0960258509990110

Ninoles, R., Planes, D., Arjona, P., Ruiz-Pastor, C., Chazarra, R., Renard, J., Bueso, E., Forment, J., Serrano, R., Kranner, I., Roach, T., & Gadea, J. (2022). Comparative analysis of wild-type accessions reveals novel determinants of Arabidopsis seed longevity. PLANT CELL AND ENVIRONMENT, 45(9), 2708–2728. 10.1111/pce.14374

Peres, S. (2016). Saving the gene pool for the future: Seed banks as archives. Studies in History and Philosophy of Science Part C: Studies in History and Philosophy of Biological and Biomedical Sciences, 55, 96–104. 10.1016/j.shpsc.2015.09.002

Pritchard, H., & Dickie, J. (2003). Predicting Seed Longevity: The use and abuse of seed viability equations (pp. 653–721).

Probert, R. J., Daws, M. I., & Hay, F. R. (2009). Ecological correlates of ex situ seed longevity: A comparative study on 195 species. Annals of Botany, 104(1), 57–69. 10.1093/aob/mcp082

Rahman, S. M. A., & Ellis, R. H. (2019). Seed quality in rice is most sensitive to drought and high temperature in early seed development. Seed Science Research, 29(4), 238–249. 10.1017/S0960258519000217

Raquid, R., Kohli, A., Reinke, R., Dionisio-Sese, M., Kwak, J., Chebotarov, D., Mo, Y., & Lee, J.-S. (2021). Genetic factors enhancing seed longevity in tropical japonica rice. Current Plant Biology, 26, 100196. 10.1016/j.cpb.2021.100196

Renard, J., Niñoles, R., Martínez-Almonacid, I., Gayubas, B., Mateos-Fernández, R., Bissoli, G., Bueso, E., Serrano, R., & Gadea, J. (2020). Identification of novel seed longevity genes related to oxidative stress and seed coat by genome-wide association studies and reverse genetics. *Plant*, Cell & Environment, 43(10), 2523–2539. 10.1111/pce.13822

Sano, N., Rajjou, L., North, H. M., Debeaujon, I., Marion-Poll, A., & Seo, M. (2016). Staying Alive: Molecular Aspects of Seed Longevity. Plant and Cell Physiology, 57(4), 660–674. 10.1093/pcp/pcv186

Satyanti, A., Nicotra, A. B., Merkling, T., & Guja, L. K. (2018). Seed mass and elevation explain variation in seed longevity of Australian alpine species. Seed Science Research, 28(4), 319–331. 10.1017/S0960258518000090

Schutte, B. J., Regnier, E. E., & Harrison, S. K. (2008). The association between seed size and seed longevity among maternal families in Ambrosia trifida L. populations. Seed Science Research, 18(4), 201–211. 10.1017/S0960258508082974

Sinniah, U. R., Ellis, R. H., & John, P. (1998). Irrigation and Seed Quality Development in Rapid-cycling Brassica: Seed Germination and Longevity. Annals of Botany, 82(3), 309–314. 10.1006/anbo.1998.0748

Society for Ecological Restoration, International Network for Seed Based Restoration, & Royal Botanical Gardens Kew. (2023, February). Seed Information Database (SID). https://ser-sid.org/

Walters, C., Wheeler, L. M., & Grotenhuis, J. M. (2005a). Longevity of seeds stored in a genebank: Species characteristics. Seed Science Research, 15(1), 1–20. 10.1079/SSR2004195

Walters, C., Wheeler, L. M., & Grotenhuis, J. M. (2005b). Longevity of seeds stored in a genebank: Species characteristics. Seed Science Research, 15(1), 1–20. 10.1079/SSR2004195

Wambugu, P. W., Nyamongo, D. O., & Kirwa, E. C. (2023). Role of Seed Banks in Supporting Ecosystem and Biodiversity Conservation and Restoration. Diversity, 15(8), Article 8. 10.3390/d15080896

White, F. J., Hay, F. R., Abeli, T., & Mondoni, A. (2023). Two decades of climate change alters seed longevity in an alpine herb: Implications for ex situ seed conservation. Alpine Botany, 133(1), 11–20. 10.1007/s00035-022-00289-8

Wolkis, D., Carta, A., Rezaei, S., & Hay, F. R. (2025). Seed longevity: Analysing post-storage germination data in R to fit the viability equation. Seed Science Research, 1–8. 10.1017/S0960258524000291

Zani, D., & Müller, J. V. (2017). Climatic control of seed longevity of *Silene* during the post-zygotic phase: Do seeds from warm, dry climates possess higher maturity and desiccation tolerance than seeds from cold, wet climates? Ecological Research, 32(6), 983–994. 10.1007/s11284-017-1508-6

Zuur, A. F., Ieno, E. N., & Elphick, C. S. (2010). A protocol for data exploration to avoid common statistical problems. Methods in Ecology and Evolution, 1(1), 3–14. 10.1111/j.2041-210X.2009.00001.x

